# Effects of Renin-Angiotensin Inhibition on ACE2 and TMPRSS2 Expression: Insights into COVID-19

**DOI:** 10.1101/2020.06.08.137331

**Authors:** Congqing Wu, Dien Ye, Adam E. Mullick, Zhenyu Li, A.H. Jan Danser, Alan Daugherty, Hong S. Lu

## Abstract

Angiotensin-converting enzyme 2 (ACE2), a component of the renin-angiotensin system, is a receptor for SARS-CoV-2, the virus that causes COVID-19. To determine whether the renin-angiotensin inhibition regulates ACE2 expression, either enalapril (an angiotensin-converting enzyme inhibitor) or losartan (an AT1 receptor blocker) was infused subcutaneously to male C57BL/6J mice for two weeks. Neither enalapril nor losartan changed abundance of ACE2 mRNA in lung, ileum, kidney, and heart. Viral entry also depends on transmembrane protease serine 2 (TMPRSS2) to prime the S protein. TMPRSS2 mRNA was abundant in lungs and ileum, modest in kidney, but barely detectable in heart. TMPRSS2 mRNA abundance was not altered by either enalapril or losartan in any of the 4 tissues. Next, we determined whether depletion of angiotensinogen (AGT), the unique substrate of the renin-angiotensin system, changes ACE2 and TMPRSS2 mRNA abundance. AGT antisense oligonucleotides (ASO) were injected subcutaneously to male C57BL/6J mice for 3 weeks. Abundance of ACE2 mRNA was unchanged in any of the 4 tissues, but TMPRSS2 mRNA was significantly decreased in lungs. Our data support that the renin-angiotensin inhibition does not regulate ACE2 and hence are not likely to increase risk for COVID-19.

## Results and Discussion

Angiotensin-converting enzyme 2 (ACE2) degrades angiotensin (Ang)I and II, and additionally is a cellular receptor for SARS-CoV-2, the virus that causes COVID-19. Viral entry into host cells occurs through binding of the viral spike (S) protein and ACE2.^1^ Preclinical data suggest that renin-angiotensin system (RAS) blockers upregulate ACE2.^2,3^ As a consequence, RAS blockers have been suggested to increase the risk of developing a severe and fatal SARS-CoV-2 infection. However, recent large retrospective studies strongly argue against this hypothesis and rather suggest that RAS blockers might be protective in such patients.^4^ Since the findings on RAS blocker-induced ACE2 upregulation are inconsistent, and differed not only per type of RAS inhibitors (ACE inhibitors versus angiotensin receptor blockers [ARB]),^3^ between blockers of a certain type (i.e., between various ARBs^5^), but also per organ, and required high doses, one further option is that this ACE2 upregulation is not the unavoidable consequence of RAS suppression, but rather reflects the non-specific effects of a certain RAS blocker when applied at a high dose. Applying antisense oligonucleotides (ASO) as a tool to suppress the RAS would circumvent the latter. In the present study we therefore compared the effects of an ACE inhibitor (enalapril) and an ARB (losartan), infused subcutaneously to male C57BL/6J mice via mini osmotic pumps, with those of subcutaneously injected angiotensinogen (AGT) antisense oligonucleotides on tissue ACE2.

After 14 days of infusion, both enalapril and losartan increased plasma renin >100-fold, confirming effective RAS inhibition (**Figure A**). ACE2 mRNA abundance was determined by quantitative PCR (qPCR) in lung, ileum, kidney, and heart tissues. Neither enalapril nor losartan changed the abundance of ACE2 mRNA in any of the tissues (**Figure B**). Ferrario et al.^3^ reported that administration of lisinopril (an ACE inhibitor) or losartan for 12 days increases ACE2 mRNA abundance approximately 3 - 5 old in male rat heart. It is worth noting that ACE2 mRNA is much less abundant in heart compared with lung, ileum, and kidney (data not shown).

Viral entry also depends on transmembrane protease serine 2 (TMPRSS2) to prime S protein.^1^ TMPRSS2 mRNA was highly abundant in lung and ileum, moderately in kidney, while barely detectable in heart (data not shown). Thus, ACE2 and TMPRSS2 are co-expressed most abundantly in lung and ileum, consistent with their roles in ARS- CoV-2 infection. As with ACE2, TMPRSS2 mRNA abundance was not altered by either enalapril or losartan (**Figure C**).

**Figure.**
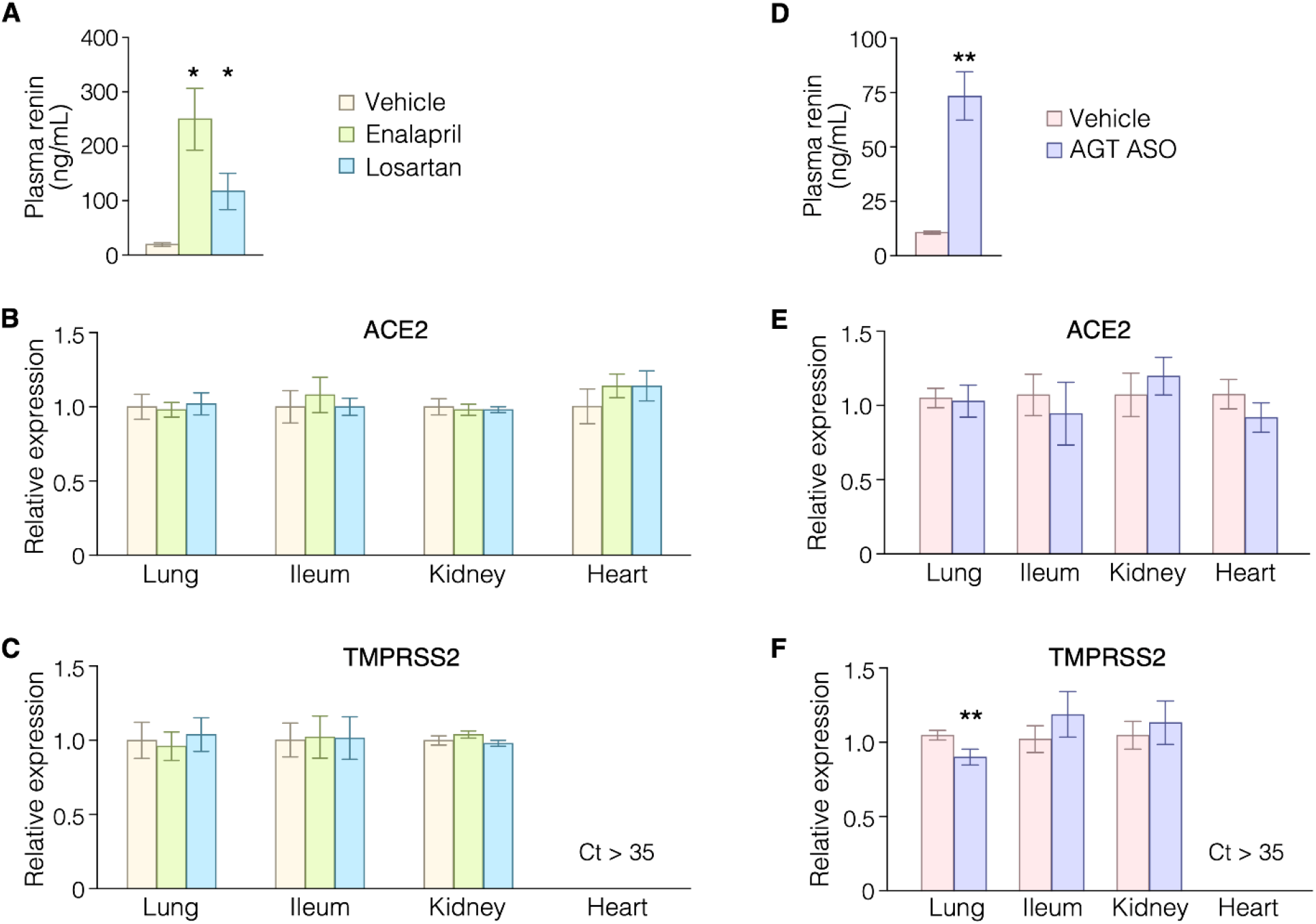
Effects of enalapril, losartan, and AGT ASO on ACE2 and TMPRSS2 mRNA abundance. **A-C.** Male C57BL/6J mice were infused subcutaneously with either enalapril (3 mg/kg/day) or losartan (15 mg/kg/day) for 14 days. **D-F.** Male C57BL/6J mice were administered AGT ASO (80 mg/kg at day 1 and 4, 40 mg/kg at day 8 and 15) via subcutaneous injection. Plasma renin concentrations (**A** and **D**) were measured using an ELISA kit. (**B, C, E, F**) ACE2 and TMPRSS2 mRNA abundance were quantified by qPCR with TaqMan probes (ID: Mm01159003_m1 and ID: Mm00443687_m1, respectively) and normalized to the geomean of three reference genes: ACTB, GAPDH, and PPIA. Genes with a cycle threshold (Ct) greater than 35 were considered undetectable. Error bars denote SEM; n = 5 for all experimental groups. * P < 0.001 versus vehicle, one-way ANOVA with Holm-Sidak method. ** P < 0.05 versus vehicle, Student’s *t*-test.

Next, we determined whether depletion of AGT, the unique substrate of the RAS, changes ACE2 and TMPRSS2 mRNA abundance. After 21 days of AGT depletion, elevated plasma renin levels supported RAS blockade (**Figure D**). However, abundance of ACE2 mRNA was unchanged in all tissues (**Figure E**). Interestingly, AGT ASO significantly decreased TMPRSS2 mRNA abundance in lungs (**Figure F**).

In summary, RAS inhibition did not affect mRNA abundance of ACE2 in male C57BL/6J mice administered enalapril, losartan, or AGT ASO. AGT ASO reduces TMPRSS2 mRNA expression in lungs, which is potentially protective against viral entry. These data support that RAS inhibition per se does not regulate ACE2 and hence is unlikely to increase the risk for COVID-19. In agreement with this concept, Sama et al.^6^ recently were unable to detect changes in circulating ACE2 in patients taking RAS inhibitors.

